# Organization and Dynamics of Crosslinked Actin Filaments in Confined Environments

**DOI:** 10.1101/2022.07.14.500066

**Authors:** Oghosa H. Akenuwa, Steven M. Abel

## Abstract

The organization of the actin cytoskeleton is impacted by the interplay between physical confinement, features of crosslinking proteins, and deformations of semiflexible actin filaments. Some crosslinking proteins preferentially bind filaments in parallel, while others bind more indiscriminately. However, a quantitative understanding of how the mode of binding influences the assembly of actin networks in confined environments is lacking. Here we employ coarse-grained computer simulations to study the dynamics and organization of semiflexible actin filaments in confined regions upon the addition of crosslinkers. We characterize how the emergent behavior is influenced by the system shape, the number and type of crosslinking proteins, and the length of filaments. Structures include isolated clusters of filaments, highly connected filament bundles, and networks of interconnected bundles and loops. Elongation of one dimension of the system promotes the formation of long bundles that align with the elongated axis. Dynamics are governed by rapid crosslinking into aggregates, followed by a slower change in their shape and connectivity. Crosslinking decreases the average bending energy of short or sparsely connected filaments by suppressing shape fluctuations. However, it increases the average bending energy in highly connected networks because filament bundles become deformed and small numbers of filaments exhibit long-lived, highly unfavorable configurations. Indiscriminate crosslinking promotes the formation of high-energy configurations due to the increased likelihood of unfavorable, difficult-to-relax configurations at early times. Taken together, this work demonstrates physical mechanisms by which crosslinker binding and physical confinement impact the emergent behavior of actin networks, which is relevant both in cells and in synthetic environments.

**SIGNIFICANCE:** The actin cytoskeleton is vital for intracellular transport, yet it remains challenging to understand how its organization is impacted by the interplay between physical confinement and the crosslinking of semiflexible actin filaments. In this study, we explore how the mode of crosslinker binding and the shape of the confining region impact the assembly and organization of actin filaments. The dynamics are governed by rapid crosslinking of spatially proximal filaments into aggregates, followed by slower relaxation of their shape and connectivity. Indiscriminate crosslinking promotes more highly connected networks, greater curvature of long filament bundles, and a subset of filaments in highly unfavorable configurations. The results provide insight into mechanisms influencing the cytoskeleton in cells and in reconstituted systems.

## INTRODUCTION

The actin cytoskeleton is a dynamic network of actin filaments that is essential for function and growth of eukaryotic cells. In plant cells, it serves as scaffolding for active, myosin-driven transport with implications for biological processes such as the transport of mitochondria to areas of metabolic activity (1), the transport of Golgi stacks to growing pollen tubes and root hairs (2–4), and cytoplasmic streaming (5). Despite the importance of actin networks in many cellular processes, it remains challenging to understand how their organization and dynamics are regulated by both biochemical and physical factors.

Actin filaments are often physically restricted within confined regions. For example, in plant cells, the cell wall provides rigid external confinement, and the large vacuole can occupy up to 90% of the cytoplasm, leading to quasi-2D environments between the vacuole and plasma membrane. The shape of cells vary based on type, leading to different confining shapes. Some cells, like root hairs and pollen tubes, are highly extended in one dimension and have large aspect ratios. Additionally, actin crosslinking proteins (ACPs) physically crosslink actin filaments in a reversible manner. Both reconstituted *in vitro* experiments and computer simulations have provided insight into the dynamics and organization of actin networks. However, much remains unknown about how they are impacted by physical confinement and properties of ACPs. In this work, we investigate confined regions of different shapes and the mode of binding of the crosslinker - whether restricted to crosslinking locally aligned filaments or indiscriminate in binding. Given the wide variety of crosslinking proteins and cellular shapes, this has the potential to inform understanding of actin networks in both cellular and reconstituted systems.

In cells, ACPs help to organize actin networks, contributing to the formation of functional structures such as lamellar networks, filopodial cables, asters, and contractile bundles (5–7). How ACPs bind to actin filaments influences the structure and properties of the actin network (8–11). Some ACPs, such as fascin and plant villin, crosslink actin filaments that are aligned in the same direction. Other ACPs, such as α-actinin and filamin, are promiscuous crosslinkers that crosslink filaments regardless of their relative orientation (5, 12). ACPs that crosslink filaments in parallel promote the formation of bundles containing filaments parallel to one another (5, 13, 14). Bundles of filaments can further influence the structure of the network because they have a larger bending stiffness than single filaments (13). Promiscuous crosslinkers promote the formation of meshwork networks comprised of both bundles and individual filaments (9, 15). However, most experimental work studying the effects of crosslinking proteins on actin networks has examined reconstituted bulk systems in which confinement does not play a role.

Computational studies have provided additional insight into the effect of ACPs on the organization of actin filaments. Most have used coarse-grained approaches where crosslinkers were modeled either as an implicit attractive force between filaments (16–19) or as a spring with explicit crosslinker-actin binding interactions (20–24). Recent approaches include MEDYAN (MEchanochemical DYnamics of Active Networks), a coarse-grained stochastic reaction-diffusion scheme (25), and AFINES (Active FIlament NEtwork Simulation), a hybrid kinetic Monte Carlo and Brownian dynamics method (20, 26, 27). Both methods have been used to simulate cytoskeletal networks with crosslinking proteins and molecular motors. Cyron et al. (22) is one of the only studies to consider preferred binding angles of crosslinked filaments, but its focus was constructing an equilibrium phase diagram when crosslinkers were limited to binding filaments at specific angles.

Both experimental and computational efforts have shown that confinement can modulate the organization of actin filaments even in the absence of crosslinking proteins. Experimental studies *in vitro* have examined the organization of actin filaments in confined regions such as microfabricated shallow chambers (28–30) and vesicles (9, 31–34). Theoretical studies have shown the formation of coils and loops by long semiflexible filaments in spherical cavities (35, 36). Relatively few studies have examined the effect of both confinement and crosslinking on actin networks. Combining actin with fascin in micropatterned, quasi-2D chambers produces bundles that tend to align with the longest axis of the confinement (30). Deshpande and Pfohl studied reconstituted actin networks in quasi-2D chambers (37), showing that the bundling agent and filament length strongly influence the organization of actin filaments. Short filaments form compact, isolated bundles whereas long filaments form a network of highly deformed, interconnected bundles. Koudehi et al. simulated actin networks with implicit attraction between polymers in spherical confinement (38). Their results showed the formation of loops, rings, and bundle structures depending on the average filament length and implicit attraction strength and range.

This paper investigates the combined effects of crosslinking and confinement on actin networks. We focus on two physical features that, to our knowledge, have not been studied together using computational methods: the shape of the region confining the actin filaments and whether crosslinkers bind in a restricted or indiscriminate manner. We extended the AFINES model originally developed by Freedman et al. (20) to account for different types of crosslinker binding. In the paper, we first give a brief overview of the computational methods and then explore the impact of system shape, mode of crosslinker binding, number of crosslinkers, and filament length. We show the impact on network organization and analyze the connectivity of the networks using graph-theoretic tools. We then characterize various measures of the dynamics and analyze the bending energy of filaments to gain insight into deformations of filaments. We discuss the results in the context of how system shape and the mode of crosslinker binding impact the dynamics and organization of actin networks.

## METHODS

### Computational framework

We used the AFINES model, developed by Freedman et al. (20, 26, 27), which is a coarse-grained model that uses kinetic Monte Carlo and Brownian dynamics to simulate actin filaments and crosslinkers. For this work, we extended the model to study crosslinkers with different binding properties, which were imposed as a potential energy term associated with the relative local orientation of crosslinked filaments. We provide a brief overview of the method, details of which can be found in Ref. (20).

In the AFINES model, actin filaments are modeled as semiflexible, bead-spring polymers in two dimensions. One end of the filament represents the barbed (plus) end and the other represents the pointed (minus) end. For a filament with *N* beads and *N* − 1 links, the potential energy is given by 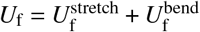, where

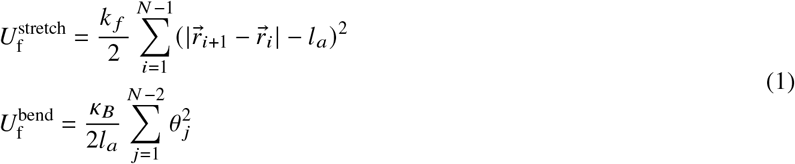

Here, 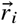 is the position of the *i*^th^ bead on the filament, *l*_*a*_ is the equilibrium spring length, k _*f*_ is the spring constant for stretching, *κ*_*B*_ is the bending modulus, and *θ* _*j*_ is the angle between the *j* ^th^ and (*j* + 1) ^th^ link. Excluded volume interactions are neglected.

Crosslinkers are treated as Hookean springs with two ends (heads) that can stochastically bind and unbind from filaments. The potential energy of a crosslinker is given by 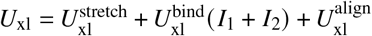, where

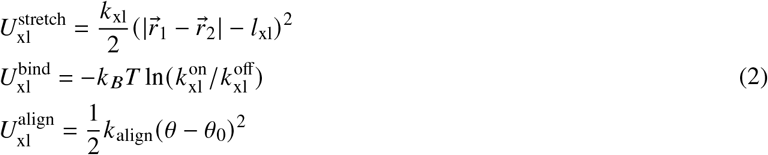

are contributions from stretching, binding to filaments, and misalignment of crosslinked filament links. Here, *l*_xl_ is the equilibrium length of the crosslinker, *k*_xl_ is its stretching stiffness, and *I*_m_ is 1 if head *m* is bound and 0 otherwise. 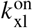 and 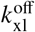 are the binding and unbinding rates respectively. For values of the parameters, we refer the reader to (26).

In this work, we introduce the angular harmonic potential, 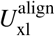, to penalize crosslinked filaments that are not locally aligned. Here, *θ* is the angle between crosslinked filament links and *θ*_0_ = 0. For *restricted crosslinkers*, which preferentially bind locally aligned filaments, *k*_align_ = 0.011 pN *µ*m. For *unrestricted crosslinkers*, which bind indiscriminately with respect to filament orientation, *k*_align_ = 0. Crosslinkers impose forces on filaments when they are concurrently bound to two filaments. This force acts on the actin beads of the filament links and is calculated in accordance with standard methods (39). When filament links *i* and *j* are crosslinked, the force due to the alignment potential acts on each of the actin beads (*a*) associated with their endpoints and is given by

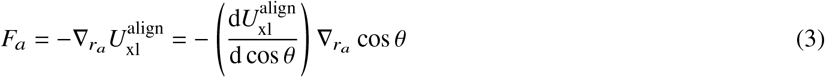

where *r*_*a*_ denotes the position of the actin bead, cos 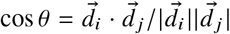, and 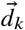 is the displacement vector of filament link *k*. Additionally, when both crosslinker heads are bound to filaments, tensile forces due to compression or stretching of the crosslinkers are propagated to the filament beads using a lever rule (24).

Dynamics are governed by a kinetic Monte Carlo arm to update the binding and unbinding of crosslinkers and a Brownian dynamics arm to update the positions of particles in the system. The system is updated at discrete times with the interval Δ*t* = 10^−4^ s. Given a state of the system, each unbound crosslinker head binds to accessible filament links with probability 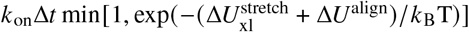. Each bound head unbinds with probability 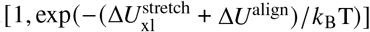. Then, given the updated state of the system, positions of the filament beads and the crosslinker heads are updated using overdamped Langevin dynamics, propagating the time forward by Δ*t*. The process is then repeated.

### Parameters studied

We considered filament lengths of *L* = 10, 5, and 2 *µ*m. The number of filaments (*N* _*f*_) was varied to keep the total filament length constant across all simulations (*N* _*f*_ *L* = 1000 *µ*m). The number of crosslinkers was varied, with *N*_*c*_ = 206, 412, 825, 1650, and 3300. Two confinement shapes were examined: a square box of size 20 *µ*m × 20 *µ*m and a rectangular box of size 40 *µ*m × 10 *µ*m.

All simulations were initialized by randomly distributing filaments in the system and allowing them to equilibrate without crosslinkers for 400 s. Unbound crosslinkers were then added uniformly at random and the simulations were continued for an additional 400 s. Reflective wall boundary conditions were imposed to simulate confinement of the actin network. For each simulation condition, 3 independent simulation trajectories were generated and analyzed.

## RESULTS AND DISCUSSION

In our simulations, we systematically varied the length of actin filaments, the number and type of crosslinkers, and the shape of confinement. To illustrate typical behavior, Fig. 1 shows snapshots taken from a trajectory with 100 filaments (*L* = 10 *µ*m) and 1650 crosslinkers (unrestricted binding, *k*_align_ = 0). A corresponding video is available (Movie S1). The trajectory exhibits an initial regime in which crosslinkers rapidly bind to filaments, crosslinking those initially in close proximity (compare *t* = 0 and 10 s). This is followed by a regime of slower relaxation in which individual filaments and bundles of filaments rearrange and coalesce, forming larger, well-defined bundles of filaments. After 400 s, the network is characterized by highly connected filaments that form several large bundles, which are curved and form loops in the network structure. A relatively small number of filaments are only partially crosslinked to the large bundles.

**Figure 1:**
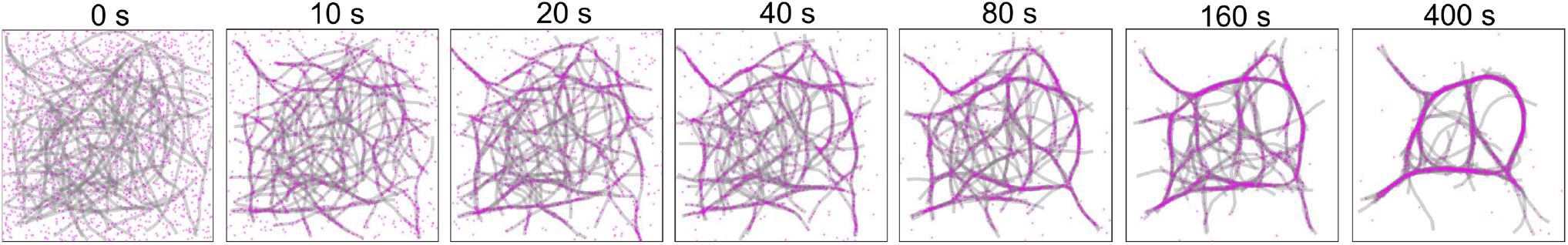
Snapshots showing the time evolution of a network with 10 *µ*m filaments and 1650 unrestricted crosslinkers in a 20 *µ*m × 20 *µ*m domain. Filaments are shown in grey and crosslinkers in fuchsia. A corresponding video is available (Movie S1).

### Network organization and filament bundling

#### Square confinement

Figure 2 shows snapshots taken from simulation trajectories in square confinement 400 s after introducing crosslinkers. The snapshots illustrate filament configurations obtained by varying filament length, the number of crosslinkers, and the alignment potential. Bundles consisting of many filaments are highlighted by the large number of crosslinkers associated with them. Figure S1 shows an alternative depiction of filaments that is analogous to an imaging experiment in which fluorophores are uniformly distributed along filaments. This highlights the local density of filaments.

**Figure 2:**
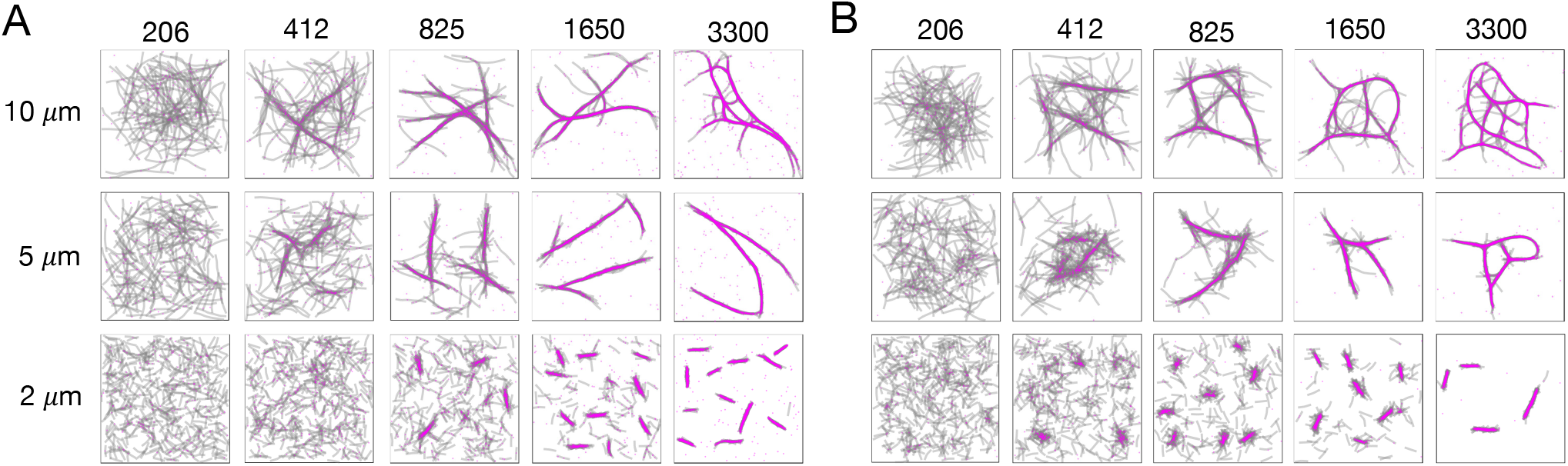
Snapshots of networks at 400 s in square confinement (20 *µ*m × 20 *µ*m) with restricted crosslinkers (A) and unrestricted crosslinkers (B). Results are shown for different filament lengths and numbers of crosslinkers. Filaments are shown in grey and crosslinkers in fuchsia.

Generally, as the number of crosslinkers increases, there is more pronounced aggregation of filaments into well-defined bundles of filaments. For small numbers of crosslinkers (206 and 412), there are localized aggregates of filaments that form bundle-like structures with longer filaments. For intermediate numbers of crosslinkers (825), most filaments are crosslinked into a small number of bundles. For larger numbers of crosslinkers, the bundles become better defined, with fewer filaments only partially connected to the bundles.

The shortest filaments (2 *µ*m) form short, isolated aggregates at intermediate and large numbers of crosslinkers. The filaments and aggregates are short compared to the confining dimensions, so the walls have little impact on their shape or their orientation. The number of distinct aggregates is larger with restricted crosslinker binding (Fig. 2A) than with unrestricted binding (Fig. 2B). This is because restricted binding prevents nearby aggregates with different orientations from easily crosslinking; it also decreases the likelihood for “spanning” filaments to connect nearby aggregates at shorter times. Further aggregation is dependent on fluctuations that change the position and orientation of aggregates, which is a relatively slow process.

Longer filaments (5 and 10 *µ*m) are more significantly impacted by confinement because they form bundled structures with lengths that exceed those of individual filaments. In these cases, the presence of confining surfaces influences the organization of the filament assemblies and leads to curvature of individual filaments and bundles. With large numbers of crosslinkers, there is an increase in the curvature of the bundles and the emergence of loops in the network structure. These features are more pronounced with unrestricted crosslinking and emerge at a smaller number of crosslinkers for 10 *µ*m filaments compared with 5 *µ*m filaments.

#### Rectangular confinement

Figure 3 shows snapshots from simulation trajectories in rectangular confinement. The cases are directly comparable to those in Fig. 2 and demonstrate the impact of the system shape. With 5 and 10 *µ*m filaments, the rectangular domain induces strong alignment of the filaments with its long dimension when the filaments are crosslinked into bundles. The alignment is quantified in Fig. S2, which shows the distribution of angles of filament links relative to the long axis of the system.

**Figure 3:**
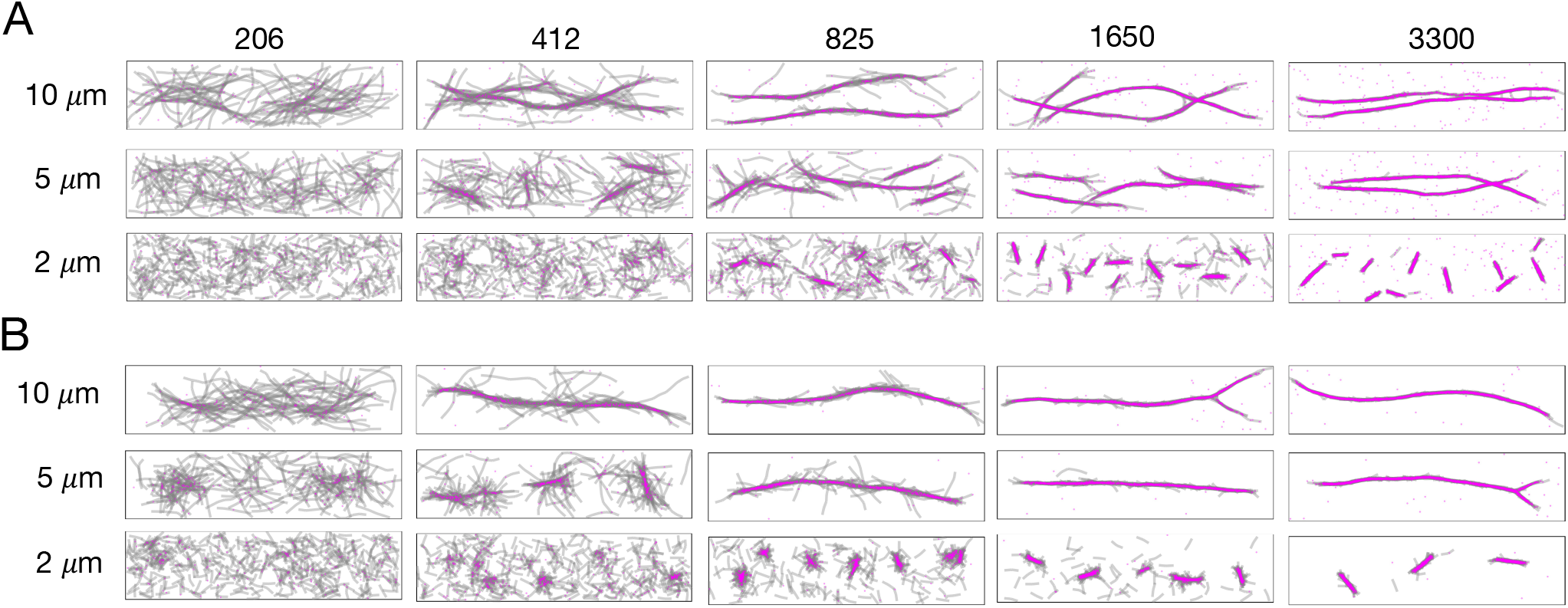
Snapshots of networks at 400 s in rectangular confinement (40 *µ*m × 10 *µ*m) with restricted crosslinkers (A) and unrestricted crosslinkers (B). Results are shown for different filament lengths and numbers of crosslinkers. Filaments are shown in grey and crosslinkers in fuchsia.

With restricted crosslinking (Fig. 3A), the filaments coalesce into two main bundles with filaments that are oriented in opposite directions. The restricted binding prevents the bundles from coalescing. The 5 *µ*m filaments require more crosslinkers to induce elongated bundles and strong orientational ordering of the filaments. With unrestricted crosslinking (Fig. 3B), small numbers of crosslinkers result in larger aggregates. Larger numbers of crosslinkers result in a single aligned bundle with a length comparable to the long dimension of the simulation box.

#### Analysis of network connectivity

To characterize connectivity of the actin networks, we represented each network as a graph in which filaments were represented as nodes. Two nodes were connected by an edge if the two corresponding filaments were crosslinked. From each graph, we parsed filaments into communities based on connectivity of the graph using the fast Newman greedy algorithm (40). The basis for the algorithm is that filaments have more connections with other filaments within their community than with filaments outside their community.

Figure 4 shows graphs depicting the connectivity of the filament networks with 10 *µ*m filaments. The nodes are colored according to their community, and corresponding snapshots are shown with the filaments colored by community. This illustrates that communities are typically associated with aggregates or bundles of filaments. Figures S3 and S4 show the graphs associated with all varied parameters (*L, N*_*c*_, and type of crosslinker).

**Figure 4:**
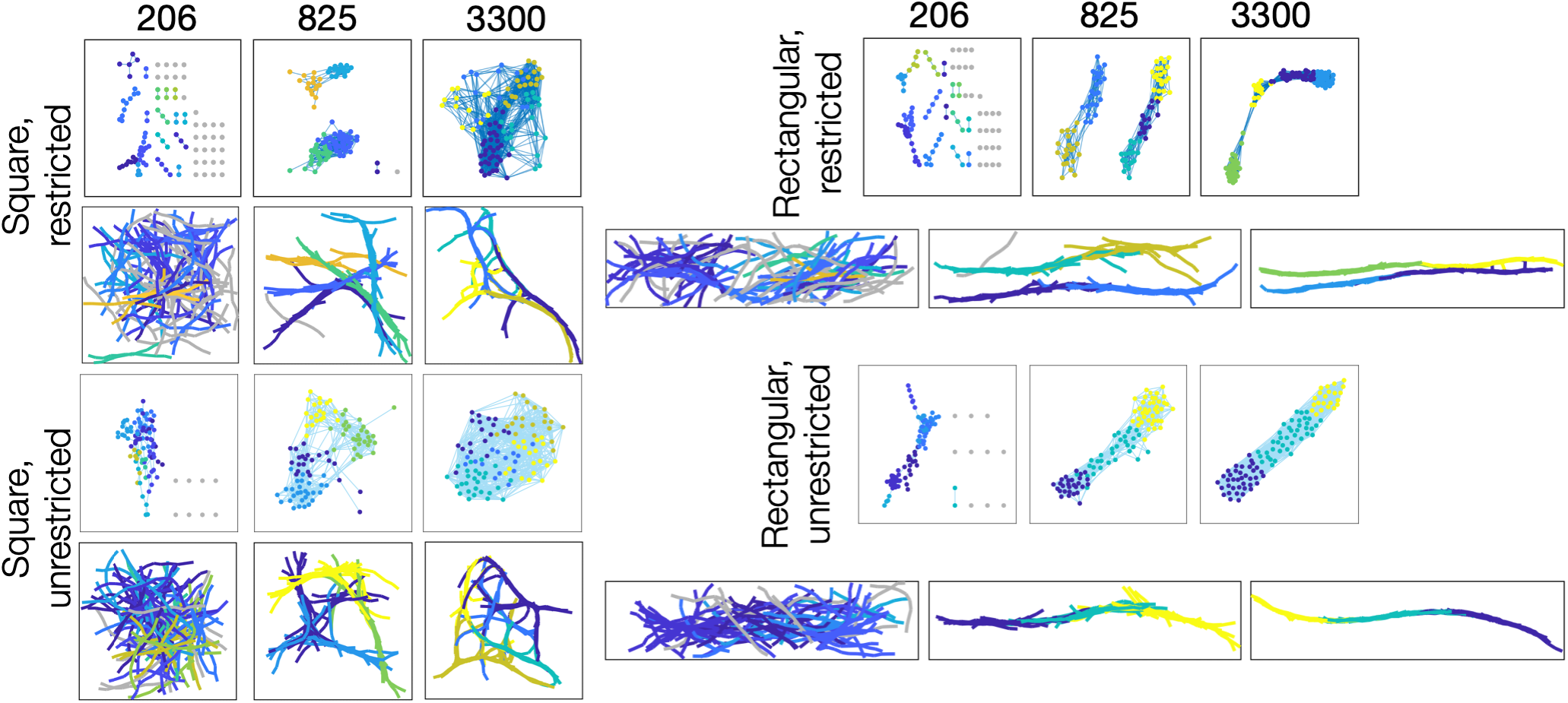
Graphs and corresponding snapshots depicting the connectivity of networks with 10 *µ*m filaments for different domain shapes and types of crosslinkers. Cases are shown with 206, 825 and 3300 crosslinkers. Each node represents a filament and each link represents crosslinking between two filaments. Nodes and corresponding filaments are colored according to the community determined based on the connectivity. Nodes and filaments that are not crosslinked are shown in gray.

With restricted crosslinkers, increasing the number of crosslinkers first leads to isolated aggregates, followed by larger bundles of filaments. With the 10 *µ*m filaments shown, large numbers of crosslinkers result in a fully connected graph, indicating that the bundles identified by community analysis are connected by filaments spanning between them. With unrestricted crosslinkers, small numbers of crosslinkers lead to more filaments being linked, and fully connected networks emerge with smaller numbers of crosslinkers. This is because the relative orientation of filaments does not impact binding, facilitating connections.

### Dynamics of aggregation and network relaxation

#### Area fraction occupied by filaments

The trajectory shown in Fig. 1 (Movie S1) illustrates that the crosslinking of filaments at short times impacts the subsequent relaxation and coarsening. Movies S2 - S4 show additional cases with the same numbers of filaments (100) and crosslinkers (1650), but that vary in crosslinker type and confinement shape. In all cases, crosslinkers lead to aggregation and bundling of filaments, but the dynamics are impacted by the type of crosslinker and the shape of confinement.

Aggregation of filaments leads to larger expanses of space without filaments. To quantify this, we computed the area fraction occupied by filaments as a function of time. The area fraction was determined by dividing the simulation box into square voxels with a side length of 0.1 *µ*m and determining the fraction of voxels containing a filament. The area fraction reported for each case is averaged over three simulation trajectories (Fig. 5). Increasing the number of crosslinkers leads to a larger decrease in the area fraction occupied by filaments over time. This is consistent with Figs. 2 and 3, with more crosslinkers promoting a higher degree of bundling. When sufficiently large numbers of crosslinkers are present, the general response is characterized by fast initial decay followed by slower decay at longer times. It is clear that many of the systems are not equilibrated after 400 s and that the area fraction would continue to decrease at longer times.

**Figure 5:**
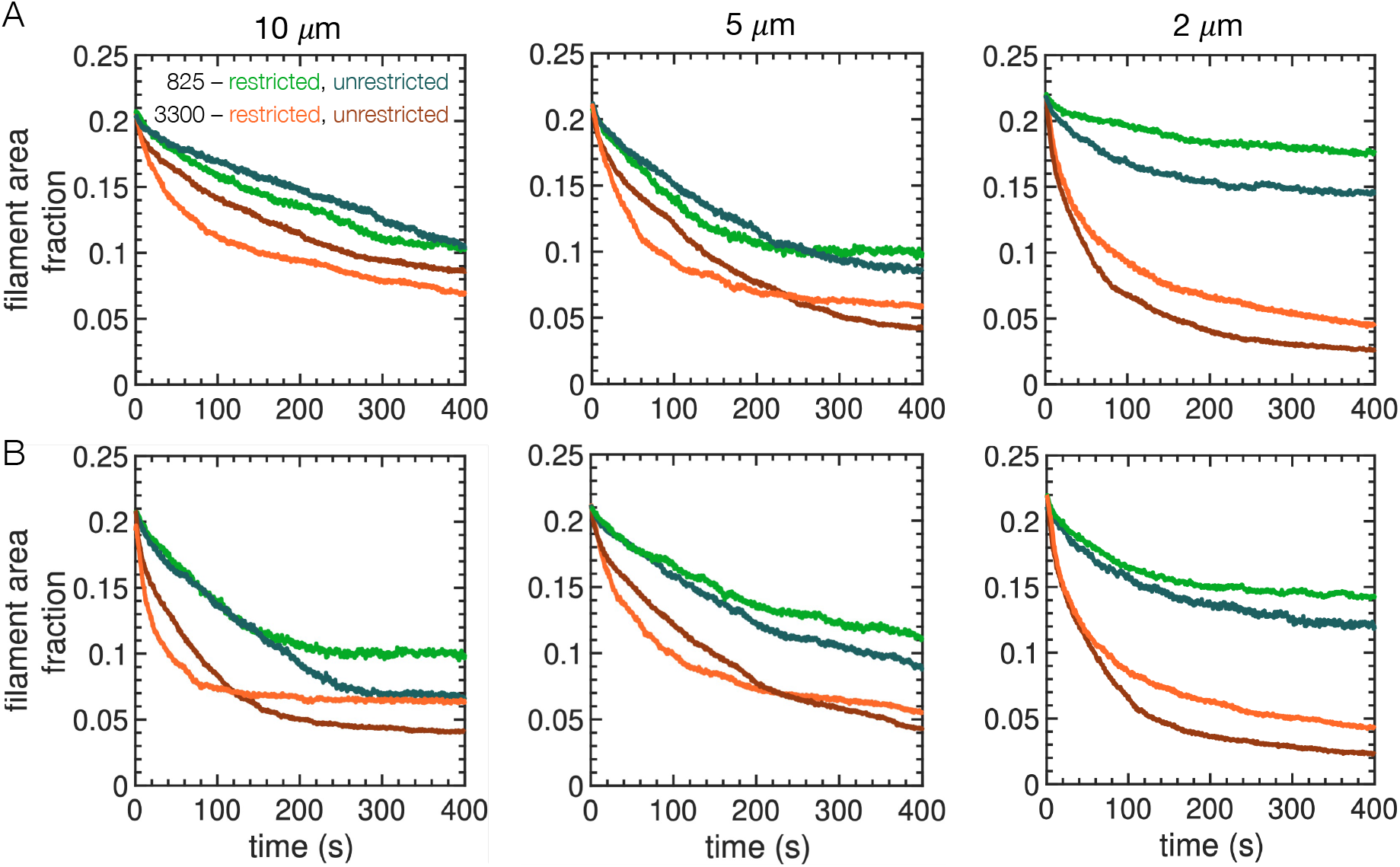
Area fraction occupied by filaments over time in square (A) and rectangular (B) domains. Curves are shown for both restricted and unrestricted crosslinkers with *N*_*c*_ = 825 and 3300. Each curve is averaged over 3 simulation trajectories.

Figure 5 also quantifies the impact of the type of crosslinker on the dynamics. With 206 crosslinkers (not shown), there is little change in the filament area fraction over time. With larger numbers of crosslinkers and 2 *µ*m filaments, unrestricted crosslinking leads to a faster and more pronounced decrease in the filament area fraction compared to restricted crosslinking.

For longer filaments, restricted crosslinkers lead to more pronounced decay of the filament area fraction at early times. However, there is a crossover point after which unrestricted crosslinkers lead to a lower filament area fraction (except for 10 *µ*m filaments in square confinement, where this would presumably occur at later times). This suggests that unrestricted crosslinking leads to frustration at early times due to the promiscuous crosslinking of filaments that is then slow to relax.

#### Connectivity of filaments

The area fraction occupied by filaments characterizes the spatial distribution of filaments, but it does not directly measure their connectivity. We therefore used the community analysis discussed above to characterize the evolving connectivity of the filaments. Figure 6 shows the number of communities as a function of time for the case with 3300 crosslinkers. After a rapid increase, the number of communities decreases over time. This corresponds to the rapid initial crosslinking of nearby filaments followed by their longer-term aggregation and rearrangement.

**Figure 6:**
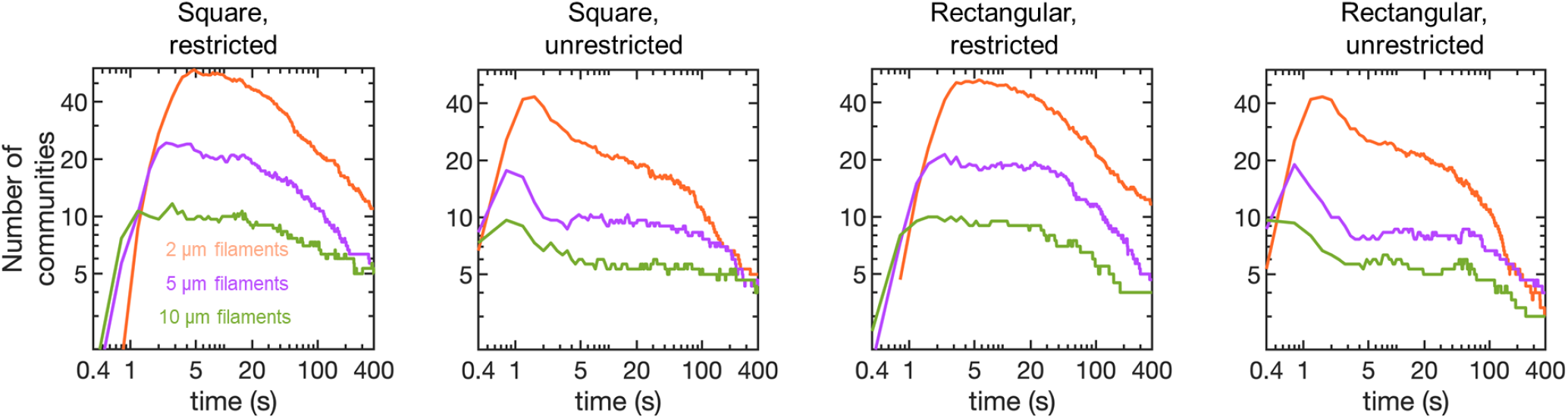
Number of communities over time. Results are shown for different filament lengths, domain shapes, and types of crosslinker. All cases have 3300 crosslinkers. Each curve is averaged over 3 simulation trajectories.

Unrestricted crosslinkers cause a rapid “overshoot” in the number of communities at short times, followed by a rapid decrease in the number of communities. In contrast, restricted crosslinkers give rise to a maximum value at a slightly later time and remain plateaued around that value for a longer time. Additionally, the maximum value is larger for restricted crosslinkers. These features are consistent with the relative ease by which unrestricted crosslinkers can bind: More filaments are directly connected at short times, leading to fewer communities. Additionally, the number of communities decays rapidly because filaments can be crosslinked and bundled regardless of orientation. In contrast, with restricted crosslinking, the initial connectivity gets “locked in” because nearby filaments oriented in different directions have to reorient before being crosslinked.

#### Physical interpretation

Differences in dynamical signatures are rooted in the interplay between crosslinker binding, filament length, and subsequent relaxation dynamics. Physically, the promiscuous binding of unrestricted crosslinkers makes it easier for nearby filaments to be crosslinked because rearrangement is not required when the filaments are not aligned. This leads to fast dynamics for short filaments because they quickly form short, bundled aggregates of nearby filaments. However, unrestricted binding can crosslink long filaments into unfavorable or hard-to-relax configurations at short times. This slows the dynamics because further relaxation relies on collective effects such as the unbinding of multiple crosslinkers or large-scale rearrangements of filaments and bundles.

### Bending energy of crosslinked filament networks

#### Average bending energy of filaments

The previous results show restricted and unrestricted crosslinkers lead to differences in the curvature of bundles (Figs. 2 and 3) and in the dynamics of aggregation (Fig. 5). These results suggest that filaments can be crosslinked in unfavorable configurations, which slows the relaxation of the network. To gain more insight into the configurations induced by crosslinking, we measured the bending energy of filaments over time as a measure of their deformations. Figure 7 shows the average energy per filament over time for various conditions. The cases with 206 and 412 crosslinkers produced similar results, so we omitted the case with 206 crosslinkers for clarity.

**Figure 7:**
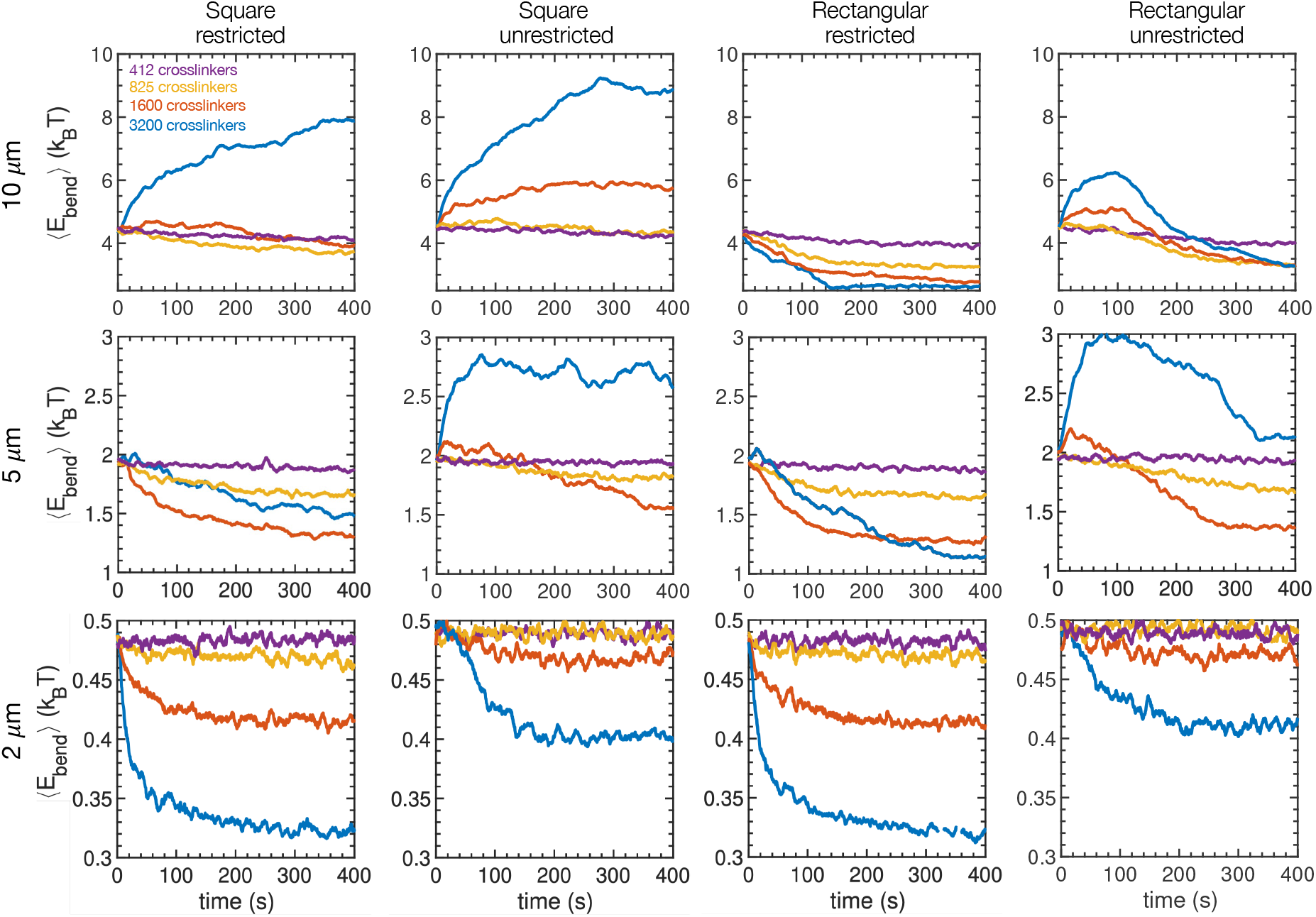
Mean filament bending energy over time. Results are shown for different filament lengths, confinement shapes, and types of crosslinker. Each curve is averaged over 3 simulation trajectories and, for clarity, is presented as a moving average with a time window of 8 s.

We begin by discussing 2 *µ*m filaments because they illustrate the influence of crosslinkers without complications introduced by strong coupling between bundles or by confinement effects. For this case (Fig. 7), the mean energy decreases with increasing number of crosslinkers. This is because the filaments form short, straight bundles. Crosslinking suppresses fluctuations in the shapes of individual filaments, thus decreasing the average bending energy. The average bending energy with restricted binding is lower than with unrestricted binding because the alignment potential promotes alignment and further suppresses shape fluctuations. The difference between square and rectangular confinement is negligible because of the short length of the aggregates.

For longer filaments, Fig. 7 reveals two key takeaways: (i) Bending energies are typically higher with unrestricted crosslinkers than with restricted, and (ii) square confinement results in higher bending energies than rectangular confinement. Small numbers of crosslinkers lead to a modest, monotonic decrease in the average bending energy. This is driven by suppressed fluctuations of crosslinked filaments. With larger numbers of crosslinkers, there is more pronounced crosslinking between different bundles, which are also long enough to be impacted by confinement. Here the behavior of the average bending energy is more complex, with increasing numbers of crosslinkers leading to less pronounced decay or even increasing energy over time (leading to nonmonotonic behavior in some cases). This behavior is more pronounced for unrestricted crosslinking and square confinement. Both of these cases promote crosslinking between different bundles of filaments and unfavorable filament configurations.

It was surprising to observe long-lived elevated bending energies in the rectangular domain (Fig. 7) because the filaments appear to form well-organized and straight bundles (Figs. 3A and B). This suggests that even though the network forms relatively straight bundles, some filaments are in highly unfavorable configurations.

#### Bending energy of individual filaments

To further explore the behavior of individual filaments, we show the bending energy of each filament over the course of a single trajectory in Fig. 8A. We focus on 10 *µ*m filaments with 3300 crosslinkers to illustrate the underlying physics. By inspection, a relatively small proportion of the filaments have markedly larger bending energies than the other filaments. To quantify this, at each timepoint we characterized “outlier filaments” as having bending energies above the upper fence of the overall distribution of bending energies (= *Q*3 + 1.5 (*Q*3 − *Q*1), where *Q*3 and *Q*1 denote the upper and lower quartile of the distribution, respectively). The energies of filaments characterized as outliers at 200 s are shown in non-gray colors.

**Figure 8:**
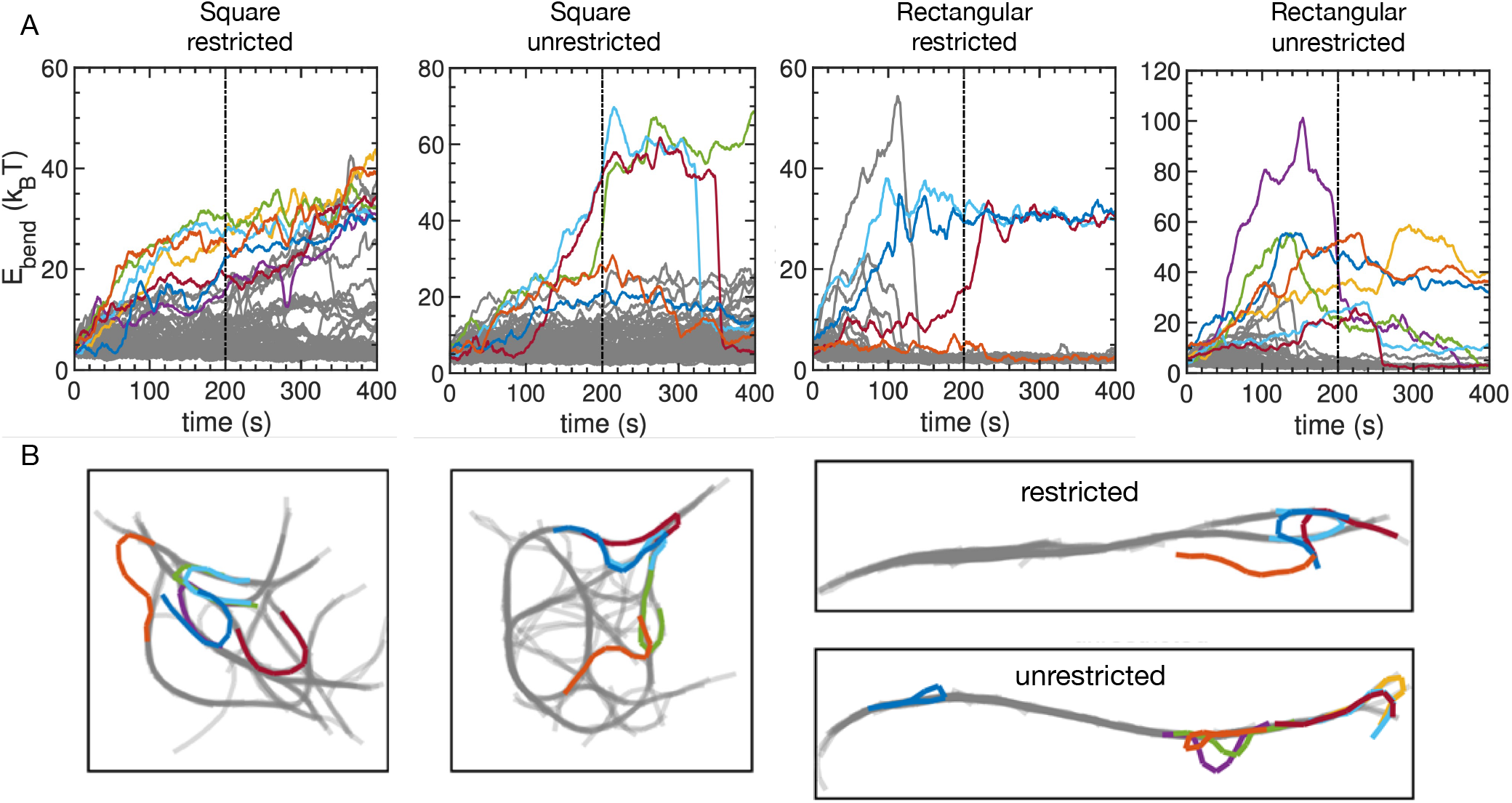
(A) Bending energies of individual filaments over the course of a simulation trajectory with 10 *µ*m filaments and 3300 crosslinkers. Filaments with bending energy that fall below the upper fence of the distribution at 200 s (vertical line) are shown in gray. The bending energy of other filaments are shown in other colors. (B) Snapshots of the same trajectories at 200 s. Filaments are shown in the same color as in (A). A corresponding video showing the full trajectory is available for each case (Movies S5 - S8).

In square confinement with restricted crosslinking (Fig. 8A), the energies of outlier filaments gradually increase over time, suggesting a slow change in the configuration of the overall network. In contrast, outliers in the other cases commonly reach sustained plateau values, suggesting long-lived, unfavorable configurations of individual filaments. Many of these rapidly switch from their high-energy state to a lower, “typical” bending energy. This indicates rapid relaxation of an individual filament from an unfavorable to a favorable configuration, which is mediated by the unbinding of crosslinkers.

Figure 8B shows snapshots taken from the trajectories in Fig. 8A. Each snapshot depicts the filament configuration at 200 s, with the outliers shown in the same color as in Fig. 8A. For the square domain with restricted crosslinking, the filaments with highest bending energy each span at least two distinct bundles and adopt a horseshoe-like shape. With unrestricted crosslinking, the outliers fall into two groups: a high-energy group with *E*_bend_ ≈ 60 *k*_B_*T* and a lower-energy group. The highest-energy filaments are distinguished by sharp, hairpin-like turns, while the lower-energy group are characterized by less severe deformations.

In rectangular confinement, restricted crosslinking leads to a small number of horseshoe-like configurations (as well as one uncrosslinked filament) that were identified as outliers. Unrestricted crosslinking leads to a variety of outliers, including hairpin-like and some that are less severely deformed. Hairpin configurations are more likely with unrestricted crosslinking because two bundles oriented in opposite directions can be crosslinked and coalesce into a single bundle. Thus, a filament that initially spans the two bundles in a horseshoe-like configuration can be forced into an even more unfavorable hairpin configuration when the bundles coalesce. This process is shown in Movies S5 - S8.

Figure 9 shows the average bending energy of filaments with and without the outlier filaments included. In square confinement, typical filaments (not including outliers) have a higher average bending energy with unrestricted crosslinkers than with restricted crosslinkers. With unrestricted crosslinking, typical bundles are more deformed, with features like loops leading to the higher average bending energy. The outliers are small in number, with few persistent hairpin configurations remaining after 400 s. This leads to a relatively small increase in the total average bending energy. With restricted crosslinkers, the typical filaments plateau at a smaller average bending energy (≈ 5.5 *k*_B_*T*) while the outlier filaments continue to increase in average bending energy. This is consistent with bundles forming early, followed by their long-time relaxation leading to an increase in energy of filaments spanning the bundles.

**Figure 9:**
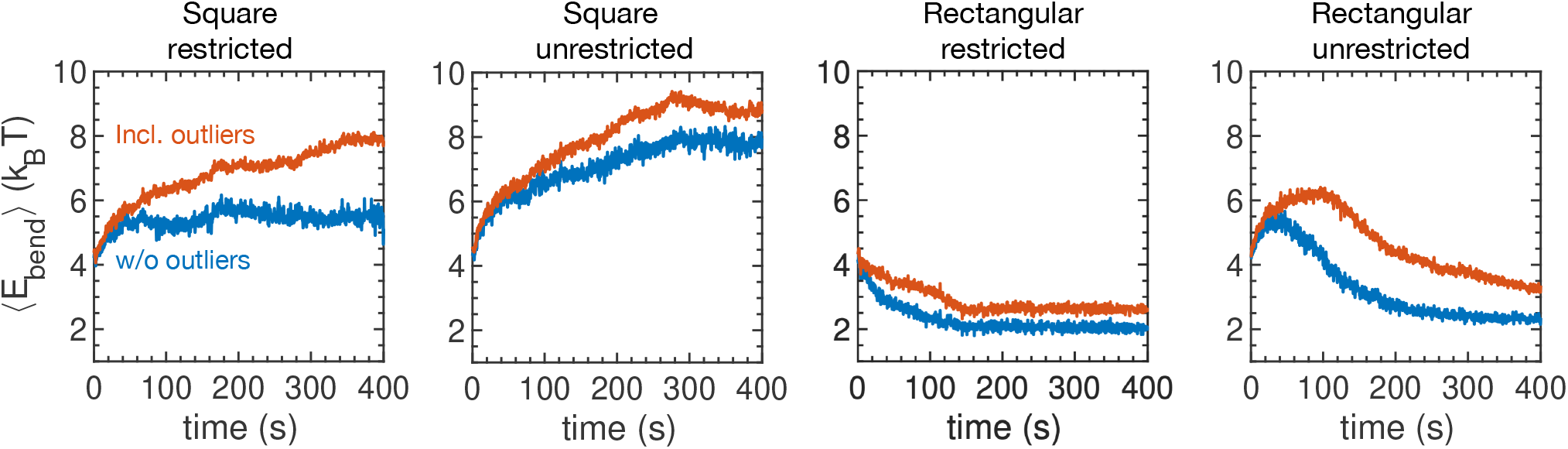
Mean filament bending energy over time for all filaments (red-orange) and with the outlier filaments at each time excluded (blue). All cases have 10 *µ*m filaments and 3300 crosslinkers. Curves are averaged over 3 trajectories.

In rectangular confinement, filaments are biased toward aligning with the long dimension of the system even before the addition of crosslinkers. This leads to less frustration when crosslinkers are introduced. With restricted crosslinkers the average bending energy without outliers rapidly decreases to a plateau, with a small number of persistent outliers leading to a modest increase in the overall bending energy. Unrestricted crosslinkers lead to a transient increase in the average bending energy without outliers, followed by a decrease to a plateau that is only slightly higher than that in the restricted case. This reflects that typical filaments have formed a single, highly crosslinked bundle spanning the system. However, the outliers lead to a larger difference than for restricted, showing the larger impact of highly deformed, hairpin-like configurations.

## CONCLUSION

Cells utilize various actin crosslinking proteins (ACPs) with different biophysical properties. This raises questions about the impact of different ACPs on the dynamics and organization of the cytoskeleton and their functional roles in cellular processes. Different ACPs also present an opportunity in reconstituted systems, where different crosslinkers could be used to control features of actin networks, with implications in applications such as artificial cells and transport in synthetic systems (41). However, fundamental questions still remain about how properties of crosslinkers and physical confinement impact cytoskeletal dynamics and organization.

Our work here demonstrates how the assembly of filaments into crosslinked networks is influenced by the mode of crosslinker binding and the shape of its physical confinement. To better understand these features, we varied the length of filaments and number of crosslinkers while focusing on two types of crosslinkers: restricted crosslinkers that preferentially crosslink locally aligned filaments and unrestricted crosslinkers that crosslink filaments without regard for relative filament orientation.

Introducing crosslinkers into a system of filaments induced aggregation and bundling of the filaments. The dynamics were characterized by a fast response at early times followed by slower changes at longer times. The fast initial decay resulted from aggregation of nearby filaments into bundles, while the slower decay reflected relaxation of the filament and bundle configurations and continued reorganization into larger bundles. With large numbers of crosslinkers and sufficiently long filaments, we observed the formation of highly connected and deformed bundles with loops and filaments spanning between bundles. This behavior was promoted by unrestricted crosslinkers and is reflected in slower aggregation (Figs. 5) and a higher average bending energy of filaments (Fig. 7). Increasing the aspect ratio of the system promoted aggregation of filaments into fewer, better-defined bundles that aligned with the long dimension of the system. It was surprising to observe elevated bending energies with relatively straight bundles in rectangular confinement. This revealed highly deformed “outlier” filaments with anomalously high bending energies. The formation of these filaments was promoted by unrestricted crosslinking and square confinement. We suggest that it would be interesting to design experiments to look for such highly deformed filaments in highly crosslinked actin networks. ≈

A number of the observations presented here are consistent with experimental results. Large numbers of crosslinkers in square confinement produced highly interconnected, deformed bundles of long filaments and short, isolated bundles of short filaments. These structures are similar to those observed in experiments in which filamin was used to crosslink actin filaments of various lengths in quasi-2D microfluidic chambers (37). Additionally, the alignment of bundles with the long axis of the system in rectangular confinement is consistent with experiments in which bundled actin filaments align with the long axis under confinement (29, 30). We also showed that large numbers of unrestricted crosslinkers caused slower aggregation of filaments when the filaments were sufficiently long. This phenomenon is similar to the dynamic arrest observed in Falzone et al., where faster growing filaments crosslinked by α-actinin generated slower bundle formation (42). In the simulations, this slower aggregation is due to unrestricted crosslinkers quickly crosslinking filaments into unfavorable conformations at early times. Combined with the semiflexible nature of filaments, this leads to frustration within the network that is difficult to relax, making further bundling more challenging. This is also consistent with accumulation and slow relaxation of stress observed in bundled networks (43, 44). Subsequent relaxation of the deformed filaments is likely controlled by thermally driven crosslinker unbinding events (44).

Our results provide insight into mechanisms contributing to the organization of actin filaments in cells and in synthetic systems. In cells, the cytoskeleton is influenced by additional factors, including forces imparted by molecular motors involved in the transport of organelles (45–48). Interesting future directions for computational studies include investigating the impact of multiple types of ACPs concurrently and studying the interplay between organelle transport and the organization of crosslinked actin networks.

## AUTHOR CONTRIBUTIONS

O.H.A. and S.M.A. designed research. O.H.A. performed research. O.H.A. and S.M.A. analyzed data and wrote the manuscript.

## ACKNOWLEDGMENTS

This work was supported by National Science Foundation grant MCB-1715794.

## SUPPORTING MATERIAL

Movie S1: 10 *µ*m filaments, 1650 unrestricted crosslinkers, square confinement

Movie S2: 10 *µ*m filaments, 1650 restricted crosslinkers, square confinement

Movie S3: 10 *µ*m filaments, 1650 unrestricted crosslinkers, rectangular confinement

Movie S4: 10 *µ*m filaments, 1650 restricted crosslinkers, rectangular confinement

Movie S5: square confinement, restricted crosslinkers (see Fig. 8)

Movie S6: square confinement, unrestricted crosslinkers (see Fig. 8)

Movie S7: rectangular confinement, restricted crosslinkers (see Fig. 8)

Movie S8: rectangular confinement, unrestricted crosslinkers (see Fig. 8)

