## Supporting Materials for "Organization and Dynamics of Crosslinked Actin Filaments in Confined Environments"

### Supporting Material

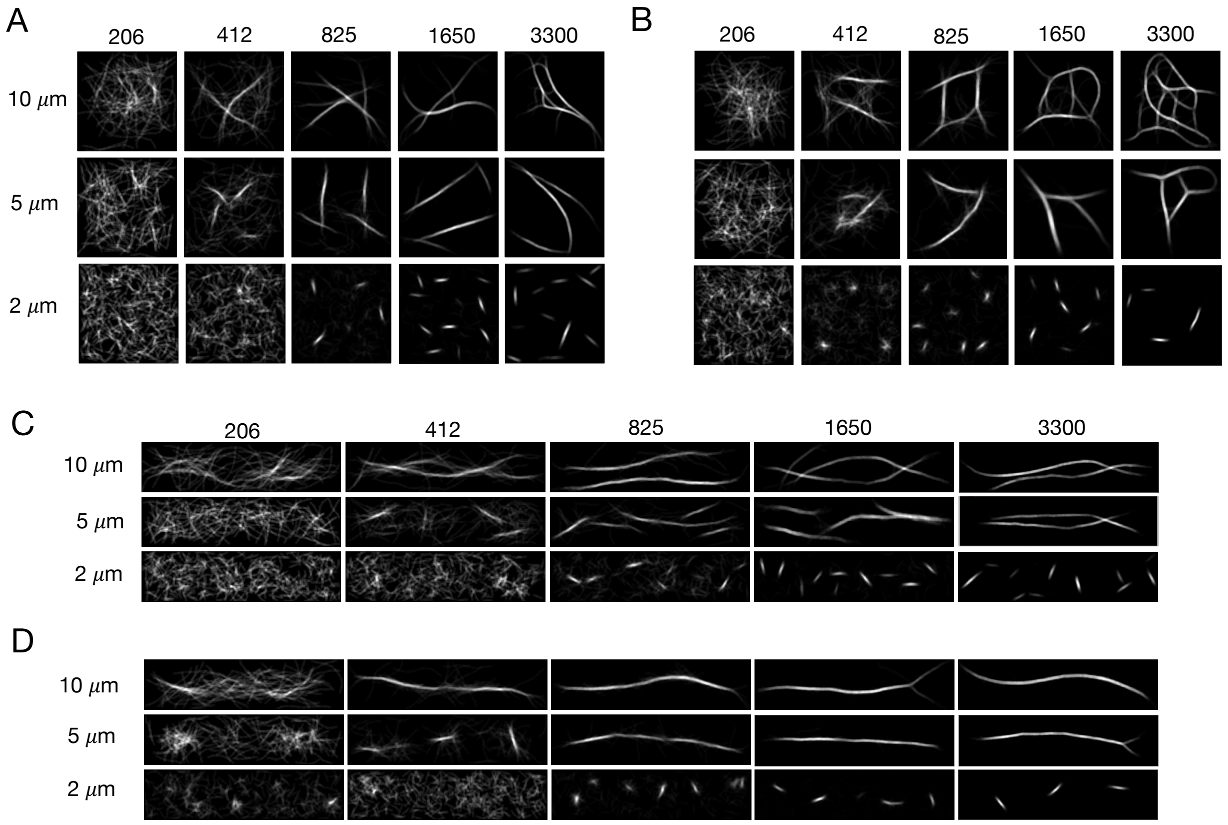


Figure S1: Snapshots of the same networks shown in Figs. 2 and 3 showing an alternative depiction of filaments that is motivated by imaging experiments in which fluorophores are distributed along filaments. The intensity is proportional to the local density of filaments. Snapshots were generated by adding point sources every 0.1 $\mu$m along each filament to mimic fluorescent markers. The system is divided into 0.1 $\mu$m $\times$ 0.1 $\mu$m voxels, and we add a constant intensity from each source for all voxels within 0.2 $\mu$m of the source. The resulting intensity map is plotted.

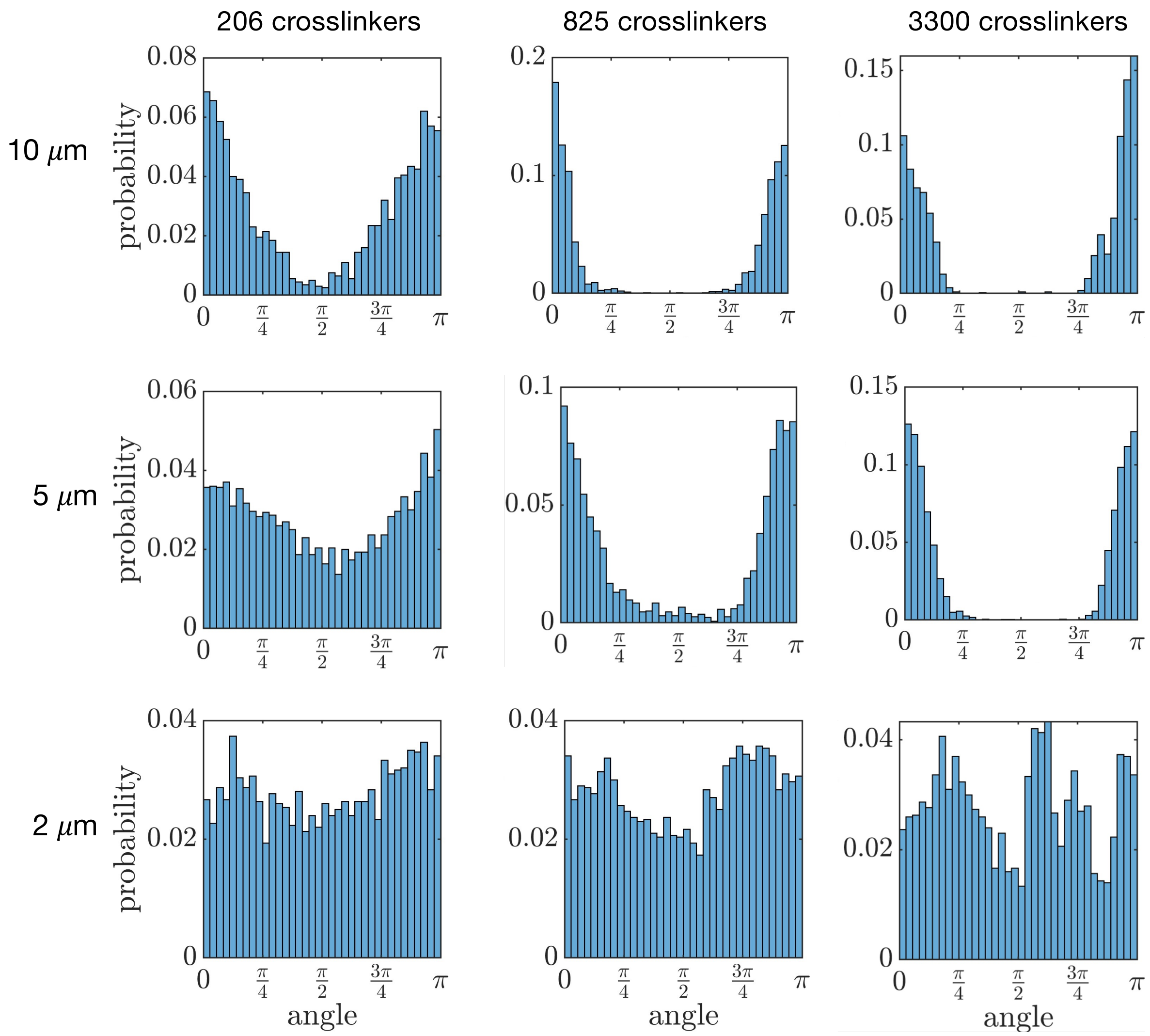


Figure S2: Distribution of angles between filament links and the +x-axis (aligned with the long dimension) for networks with restricted crosslinkers in rectangular confinement. Different filament lengths and numbers of crosslinkers are shown. Each distribution is constructed with data from 3 simulation trajectories.

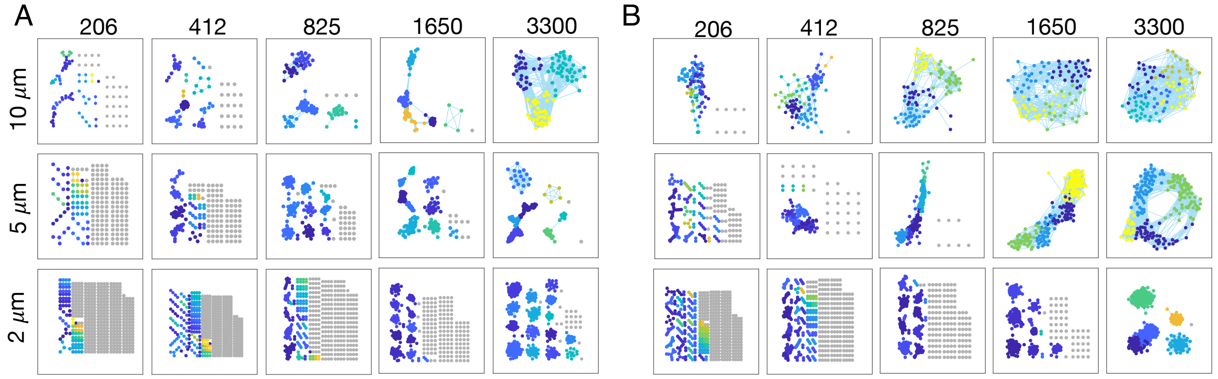


Figure S3: Graphs depicting the connectivity of networks in square confinement with restricted crosslinkers (A) and unrestricted crosslinkers (B). Results correspond to the final timepoint of a single trajectory for each case simulated. Nodes are colored according to the community determined based on the connectivity. Nodes that are not crosslinked are shown in gray.

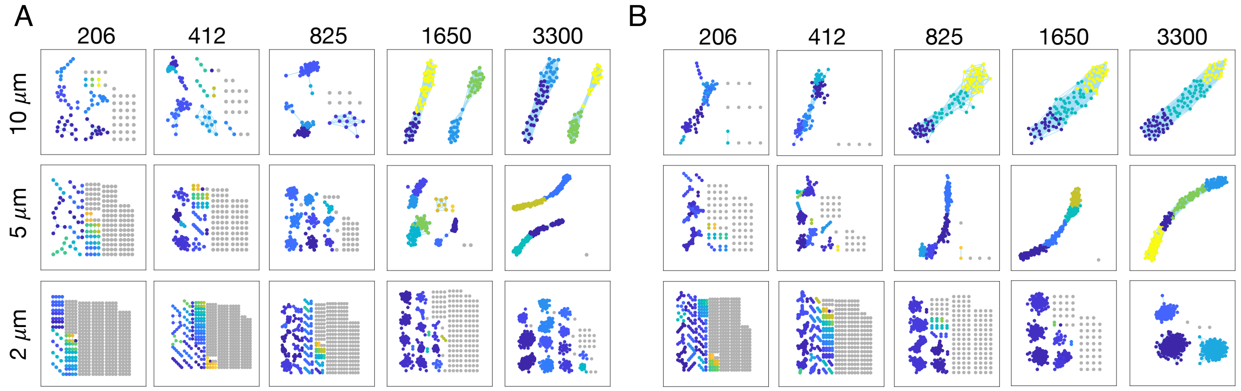


Figure S4: Graphs depicting the connectivity of networks in rectangular confinement with restricted crosslinkers (A) and unrestricted crosslinkers (B). Results correspond to the final timepoint of a single trajectory for each case simulated. Nodes are colored according to the community determined based on the connectivity. Nodes that are not crosslinked are shown in gray.
